# Novel Object Detection and Multiplexed Motion Representation in Retinal Bipolar Cells

**DOI:** 10.1101/2021.05.13.444054

**Authors:** John A. Gaynes, Samuel A. Budoff, Michael J. Grybko, Joshua B. Hunt, Alon Poleg-Polsky

## Abstract

Antagonistic interactions between the center and surround receptive field (RF) components lie at the heart of the computations performed in the visual system. Center-surround RFs are thought to enhance responses to spatial contrasts (i.e., edges), but how they contribute to motion processing is unknown. Here, we addressed this question in retinal bipolar cells, the first visual neuron with classic center-surround interactions. We found that bipolar glutamate release emphasizes objects that emerge in the RF; their responses to continuous motion are smaller, slower, and cannot be predicted by signals elicited by stationary stimuli. The alteration in signal dynamics induced by novel objects dwarfs the enhancement of spatial edges and can be explained by priming of RF surround during continuous motion. These findings echo the salience of human visual perception and demonstrate an unappreciated capacity of the center-surround architecture to facilitate novel object detection and multiplexed encoding of distinct sensory modalities.

## Introduction

The ability to detect motion begins in the retina, which contains ganglion cells dedicated to the detection of local motion^1–3^, approaching objects^4^, acceleration^5^ and the direction of movement (for a review, see^6,7^). The highly specialized computations in ganglion cells are driven and shaped by glutamate release from axonal terminals of bipolar cells (BCs), which in mice are divided into about 14-15 functional types that are tuned to different visual features^8–10^. The topographic stratification of BC axons in the inner plexiform layer (IPL) establishes some of the functional organization of visual processing in the retina: BCs that carry ON signals (depolarization to light) are found closer to the ganglion cell layer, and cells with sustained responses are segregated towards IPL borders^8,11–13^. The difference in visual processing between BCs reflects their center-surround architecture, comprised of two separate concentric regions sampling the visual signal. This RF structure is formed by direct innervation of BC dendrites by photoreceptors in their excitatory center and a combination of horizontal and amacrine cell inhibition in the antagonistic surround^9,14–16^.

Historically, motion signals in BCs have been understood as a linear combination of static responses, much like how the perception of motion is produced in movies by a rapid presentation of discrete images^17,18^. However, computations in cells with centre-surround RFs are inherently nonlinear^19–22^ and depend on the spatiotemporal RF activation pattern, which differs between moving and static stimuli. Thus, despite the abundance of the classic centre-surround RFs in the early visual system, little is known about their impact on motion processing in general and on BC activity in particular.

To examine the properties of visual processing of moving objects in BCs, we recorded the change in glutamate levels across different depths of the inner plexiform layer (IPL) and captured the release dynamics of different BC types to moving or stationary bars. We reveal significant alteration in the peak and the temporal characteristics of the glutamate responses following object motion. Additionally, our results indicate that BCs can signal the appearance of novel objects that enter the visual scene. Flashed stationary objects or stimuli that emerge from behind static occluders provoke intense discharge from all BCs, whereas continuous motion and disappearing stimuli suppress BC activation. These observations were not affected by the pharmacological blockage of amacrine cell inhibition. Accordingly, a detailed simulation of signaling in the outer retina replicates the diversity of motion responses in BCs and reveals how motion computations can be carried out at the first retinal synapse by a horizontal cell-derived inhibitory signal and influence the representation of a realistic visual input. Our results reveal a fundamental property of signal integration in center-surround RFs to identify newly appearing visual stimuli and diversify the representation of static and moving shapes.

## Results

### Glutamate responses in BCs to full-field motion are diverse and do not follow the response dynamics for stationary signals

To study the representation of moving stimuli in BCs, we used two-photon microscopy to collect light-driven glutamatergic signals in whole-mount mouse retinas expressing iGluSnFR, responding to static flashes and full-field moving bars^9,13,23^. We systematically surveyed all layers of the inner plexiform layer (IPL) with multiple scan fields; pixels with similar responses were then grouped into regions of interest (ROIs, Figs. 1a-b, s1, s2). The spatial extent of most ROIs was smaller than 50 μm, indicating sampling from a single cell or at most two functionally similar BCs (Figs. 1b, s1)^9,24^. Responses to stationary flashes were used to combine ROIs from different experiments into functional clusters^9,11,24^. The optimal separation was obtained with 5 OFF and 7 ON clusters (Figs. 1c-d); comparable to previous classifications of glutamate signals in the IPL^9,11^. Following clustering, we analyzed responses to moving bars. As expected, slower RF engagement prolonged motion response kinetics (Fig. 1d-f). Surprisingly, there was no correlation between the flash- and motion-driven rise time dynamics (Fig. 1f, left, Pearson correlation coefficient, R=-0.04).

**Fig. 1.**
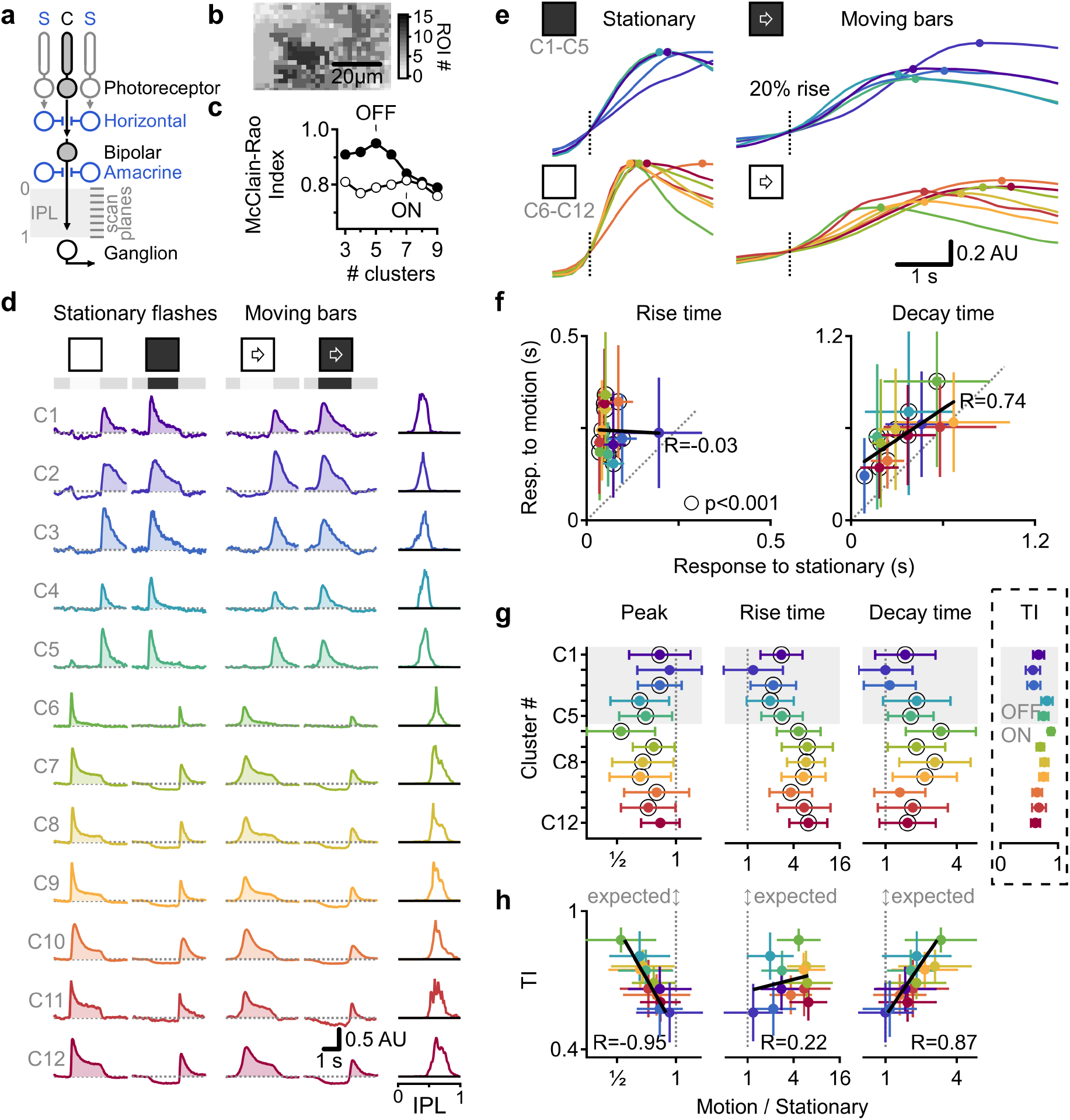
Multiplexed representation of static and moving objects in BCs. **a** Centre-surround RF structure in BCs. **b** Exemplar ROIs identified from iGluSnFR fluorescence in a single scan plane. **c** Diversity of responses to stationary flashes from 1265 ROIs suggests 12 functional clusters of glutamate release. **d R**esponses from the identified clusters, sorted by pixel depth distribution (right). **e** Focus on the rising phase of the signals. Circles indicate peaks. **f** Mean (±SD) clusters’ kinetics, linear fits in black. **g-h** IPL depth (**g**) or transiency index (**h**, inset in **g**) vs. the mean (±SD) responses ratio.

It is possible that the difference in motion processing we describe reflects the topographic stratification of BC axons with sustained responses approximating the IPL borders^8,11–13^. To assess this, we analyzed signal parameters relative to recording depth (Fig. 1g) or signal transiency index (TI, calculated from stationary response kinetics, Fig. 1g-h). We identified a clear relationship between cluster transiency to the change in the amplitude (R = - 0.95) and the decay-time (R = 0.87) of motion responses relative to the stationary signals (Fig. 1h). Similarly, the effect of motion on the peak amplitude and decay time was greatest in the central regions of the IPL, reflecting the stratification level of the transient BCs (Fig. 1g). In contrast, the change in the rise-time did not follow the transient-sustained division (Fig. 1h). Instead, we observed a gradual decrease in the motion/stationary ratio for the rise-time kinetics with increasing depth in the retina (Fig. 1g). Overall, the observed low correlation in key aspects of response shape and the distinct pattern of signal dependency on IPL depth between static and moving objects indicate a multiplexed representation of motion and stationary information in the BC population.

Notably, these observations are not an artifact of our clustering approach, as our algorithm was agnostic to motion information. We conducted several tests to rule out the possibility that the results we describe here are due to the grouping of pixels with different recruitment times during motion responses. First, at odds with the predicted effects of such pixel averaging, the degree to which motion impacted signal dynamics varied systematically between clusters, and the inter-cluster variability of responses was higher during motion (Fig. 1d). Second, the mean responses recorded for each group closely mirrored the signals recorded in individual pixels (Fig. s2). Last, neighboring regions of the retina respond sequentially to motion, and for this reason, the influence of pixel averaging should be most evident in groups with wide spatial pixel distribution. In contrast to this prediction, however, we found that the spread of each group’s pixels along the axis of motion was not correlated with the response dynamics (Fig. s3).

### Enhanced representation of novel stimuli

Previous work demonstrated that neurons could employ a simple strategy of comparing the spatial extent of center-surround recruitment to detect local spatial contrasts^19,25,26^ and diversify the representation of flashed objects^9,21^. According to the classic description of the center-surround interactions, occluders masking part of the surround enhance RF output (Fig. 2a ‘Edge’). We reasoned that responses to moving stimuli should also be sensitive to stationary edges in the RF. To explore this possibility, we presented horizontally moving bars and masked the stimulus on the left or the right halves of the display.

**Fig. 2.**
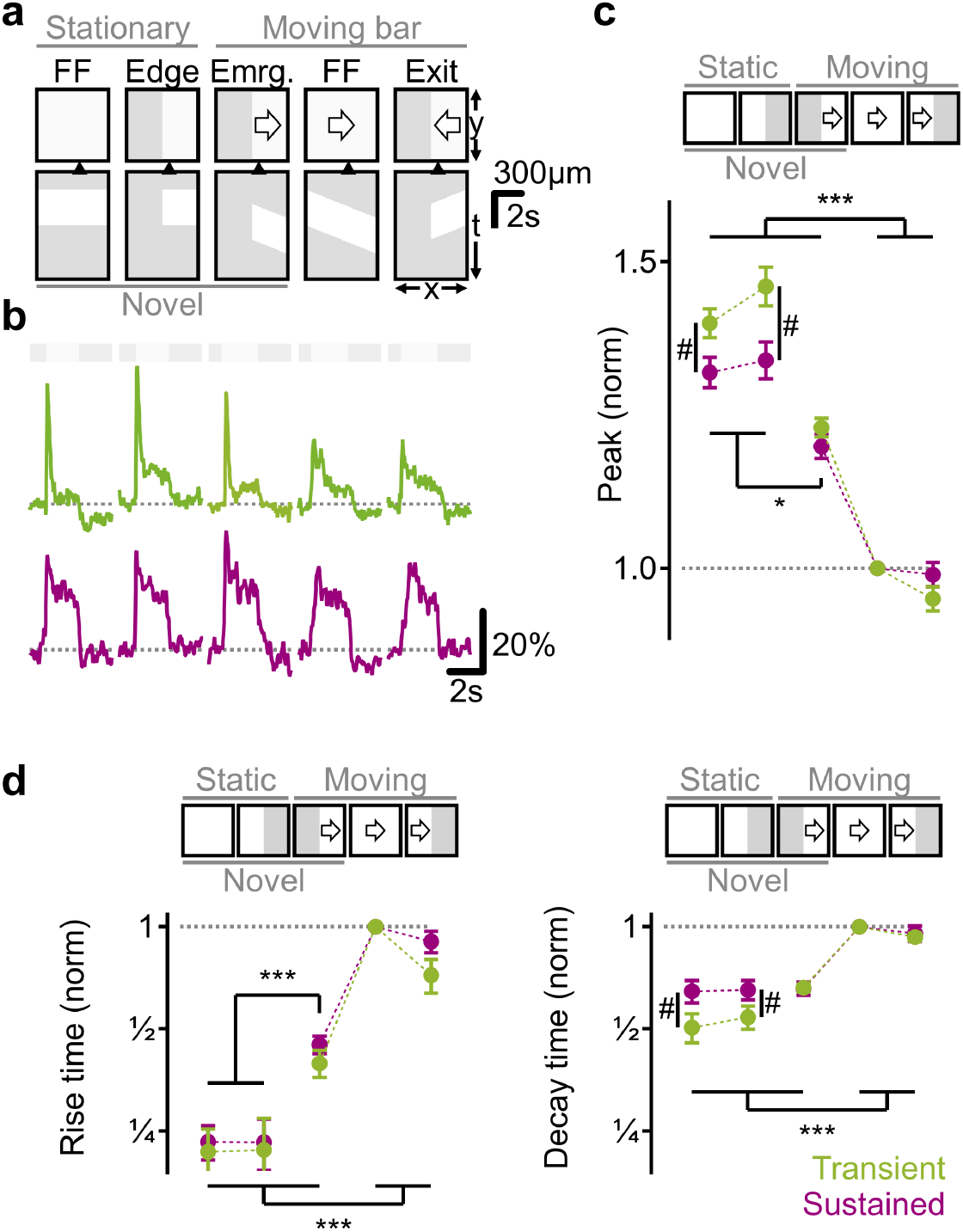
BCs release signals novel object appearance. **a** Visual protocols. For each: spatial arrangement (left) of the mask (grey)/stimulus (white) and time-space plot (right). Triangle-horizontal RF position. **b** Responses from ROIs (green, transient; red, sustained) located up to 50 μm from the mask-stimulus boundary. **c-d** Peak glutamate fluorescence (**c**), and rise and decay times (**d**) normalized by full-field motion. *p<0.05,***p<0.001 for both kinetics; #p<0.001 transient vs. sustained. Error bars-SEM.

Unexpectedly, the kinetics and the amplitude of the glutamate release were significantly faster/higher for bars emerging from the mask than for motion in the opposite direction (Fig. 2 ‘Emergence’ vs. ‘Exit’). Across all ROIs, the peak response amplitude following emerging motion was significantly higher than the signal observed during full-field motion (122±2% mean±SEM; p<0.001 vs. full-field motion, ANOVA followed by Tukey test), and the rise-time was sharpened by more than 50% (Fig. 2d). In comparison, the mean(±SEM) ratio between responses to fullfield flashes and motion in the same ROIs was 136±2.4% (p<0.001, ANOVA followed by Tukey test, n=365, Fig. 2). Thus, in terms of shape peak and temporal dynamics, the representation of emerging motion more closely resembles static flashes than continuous motion (Fig. 2).

We next separated between and compared transient (n = 182) and sustained (n = 183) ROIs to assess the effect of signal kinetics on visual processing in the presence of edges. The peak glutamate signal recorded for static flashes and emerging objects was significantly higher in ROIs with transient signals (Figs. 2c, s4), in line with the abovementioned differences in motion signaling between the transient and sustained populations (Fig. 1g-h). In contrast, the dynamics of motion exit were indistinguishable from continuous motion (Figs. 2, s4).

Compared to the classic role of center-surround in detecting spatial boundaries, we note that even though the occluding mask (when present) was identical for all protocols, we did not observe a significant effect of the masks on stationary responses (Figs. 2, s4 p>0.3 for peak/kinetics, ANOVA). We interpret this finding to indicate that RF structure in the early visual system is tuned to highlight new information and not to detect inhomogeneous spatial compositions.

### Amacrine cell inhibition is not required for novel object sensitivity and slower motion kinetics

What are the cellular components underlying motion computations in BCs? Previous work suggested that amacrine cell inhibition diversifies the representation of stationary stimuli that partially occupy the RF of BCs^9^. Correspondingly, a cocktail of 50μM SR95531, 100μM TPMPA, and 1μM Strychnine to block GABA_A_, GABA_C_ and glycine receptors (Fig. 3a)^9,24^ changed the shape of glutamate waveforms elicited by stationary flashes (Figs. 3) and of calcium transients in ganglion cells (Fig. s5). Yet, motion signals in the IPL were not affected by the inhibitory blockers (Fig. 3). Because horizontal cells can control photoreceptor output by mechanisms that do not require the release of neurotransmitters^16,27,28^, we reasoned that our pharmacological manipulation did not fully disrupt the horizontal feedback on the photoreceptors, suggesting that motion processing is performed already in the first retinal synapse.

**Fig. 3.**
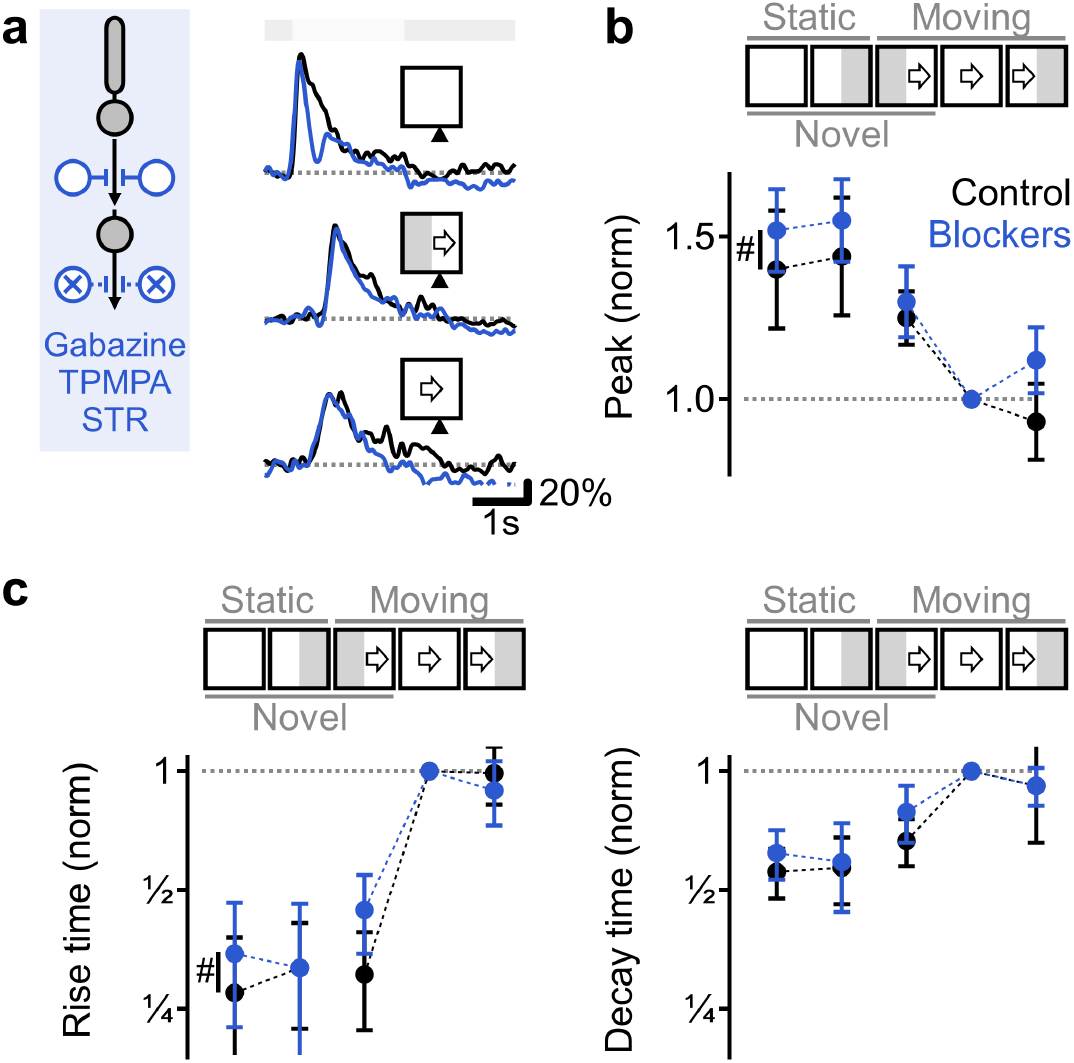
Pharmacological blockage of amacrine cell inhibition does not eliminate novel object sensitivity and slower motion response kinetics. **a** Representative glutamate responses before (black) and after (blue) blockage of amacrine cell inhibition. **b-c** Peak glutamate fluorescence (**b**), and rise and decay times (**c**) normalized by full-field motion. #p<0.001 control vs. blockers. ANOVA followed by Tukey test with Bonferroni’s correction. Error bars-SEM.

### A common horizontal cell-mediated surround can drive diverse motion responses

To test whether signal interactions in the outer retina are sufficient to explain our experimental findings, we constructed a computational model of visual processing in the outer retina and BCs (Fig. 4). We activated the model with stationary and moving bars and recorded the resulting signals in photoreceptors, horizontal cells, and BCs. The simulation revealed a possibility for a pronounced representation of emerging motion already in photoreceptors (Figs. 4), but only when horizontal cell inhibition was intact (Fig. s6). The mechanistic implementation of this outcome relied on the lag between activation times of photoreceptors and horizontal cells. At the location of object emergence, horizontal cell engagement coincided with photoreceptor activation (Figs. 4b, d, s7). Elsewhere, the initiation of horizontal cell signal preceded direct light-induced photoreceptor activation by as much as ~100 ms and correlated with diminished photoreceptor output (Fig. 4b, d, s7). While our model incorporated nonlinear interactions between cells and synaptic inputs, a similar temporal relationship in RF activation was readily observed in a linear center-surround architecture (Fig. s8). In both models, established (continuous) motion recruits the RF consecutively because moving objects encounter the surround first. RF components were engaged more synchronously by emerging stimuli and activated simultaneously by stationary flashes (Figs. 4). In general, the encoding of existing objects is accompanied by a longer temporal delay between the initial activation of the surround and subsequent center stimulation. Due to this delay, surround inhibition is more developed by the time the center is engaged by the stimulus and is, therefore, more likely to suppress responses to continuous motion.

**Fig. 4.**
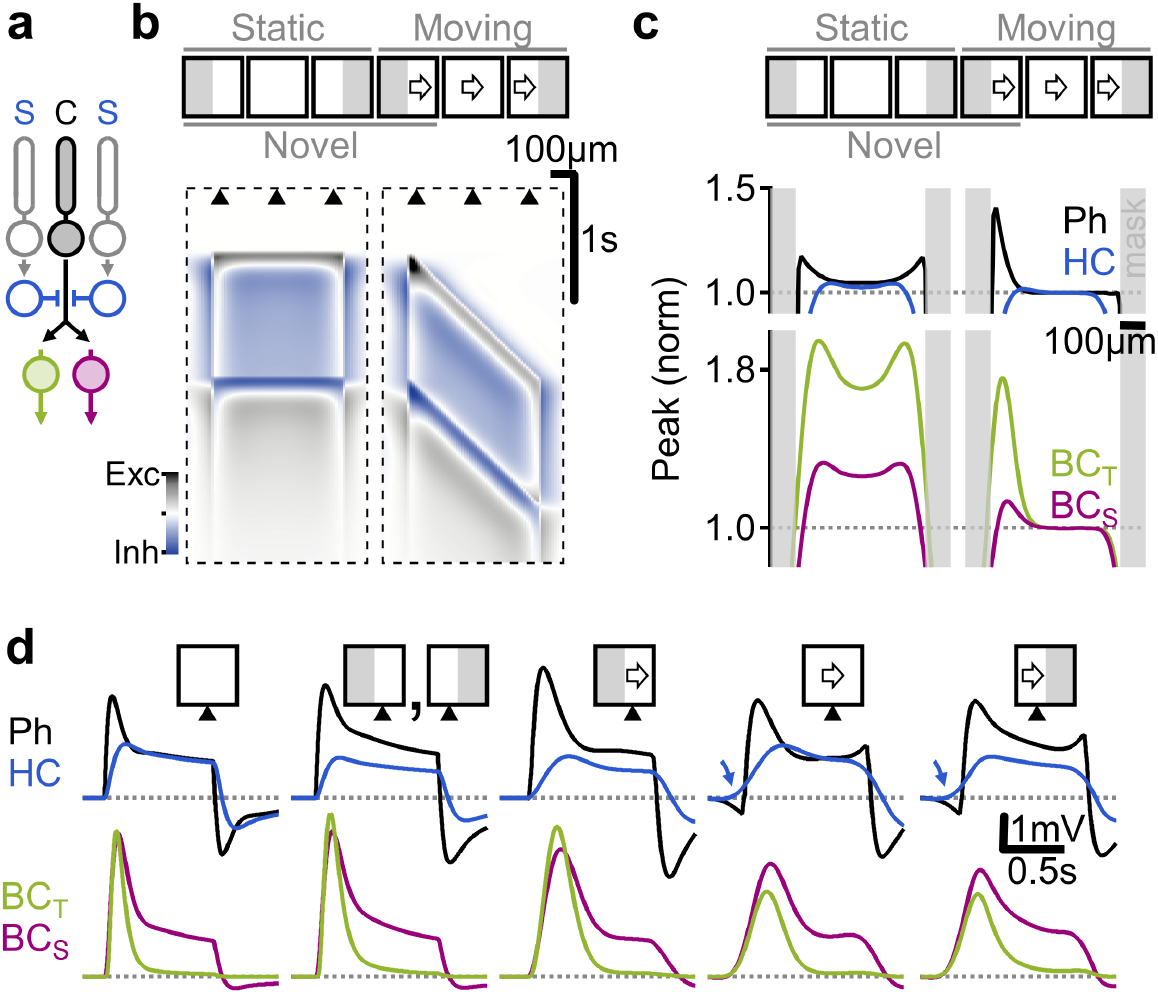
Detailed simulation of the first retinal synapse captures empirically observed novel object enhancement and population dynamics. **a** The circuit modelled in **b-d**. **b** Space-time plot showing the difference between photoreceptive and horizontal cell potentials. **c** Peak depolarization vs. spatial positions; the kinetics of the BCs (bottom) differ due to simulated BC sensitivity to photoreceptive release. Photoreceptive and horizontal cell voltages are inverted. **d** Responses from cells in positions marked in **b**. Arrows, preceding inhibition.

Focusing on the factors that influence BC response dynamics, we note that transient kinetics were mediated by faster neurotransmission, but also, unexpectedly, by elevated sensitivities to photoreceptor release (Figs. s8, s7). The higher threshold required for effectual activation increased the sensitivity of transient BCs to small fluctuations around the peak photoreceptor activity. In agreement with recent findings^22^, our model suggests that transient BCs receive a more rectified, nonlinear copy of the photoreceptor signal and predicts that such nonlinearity creates a substrate for more distinct responses to motion vs. stationary stimuli and promotes the enhancement of novel object emergence (Figs. 4c-d, s9).

### Enhanced representation of novel stimuli under natural movies requires center-surround organization

Next, we asked whether the fundamental properties of the center-surround RF architecture are sufficient to identify novel objects under realistic visual conditions. To address this question, we simulated responses from a population of linear center-surround neurons (Fig. s9) to movies showing the appearance of predators in a natural mouse habitat (Fig. 5a). Although the simulated cells lacked nonlinear signal processing mechanisms, we found these cells capable of generating a rich representation of dynamically changing scenes. Cells responding to established motion encoded the local contrast differences between the stimulus and the background (Figs. 5b-c). Comparable to our findings presented above, stimulus emergence correlated with robust responses (Fig. 5b-c, s10). Interestingly, novel motion enhancement was evident mainly at the initial site of stimulus appearance (the wing in the example shown in Fig. 5), implying a spatial focus for novel object detection spanning about 100 μm of retinal space.

**Fig. 5.**
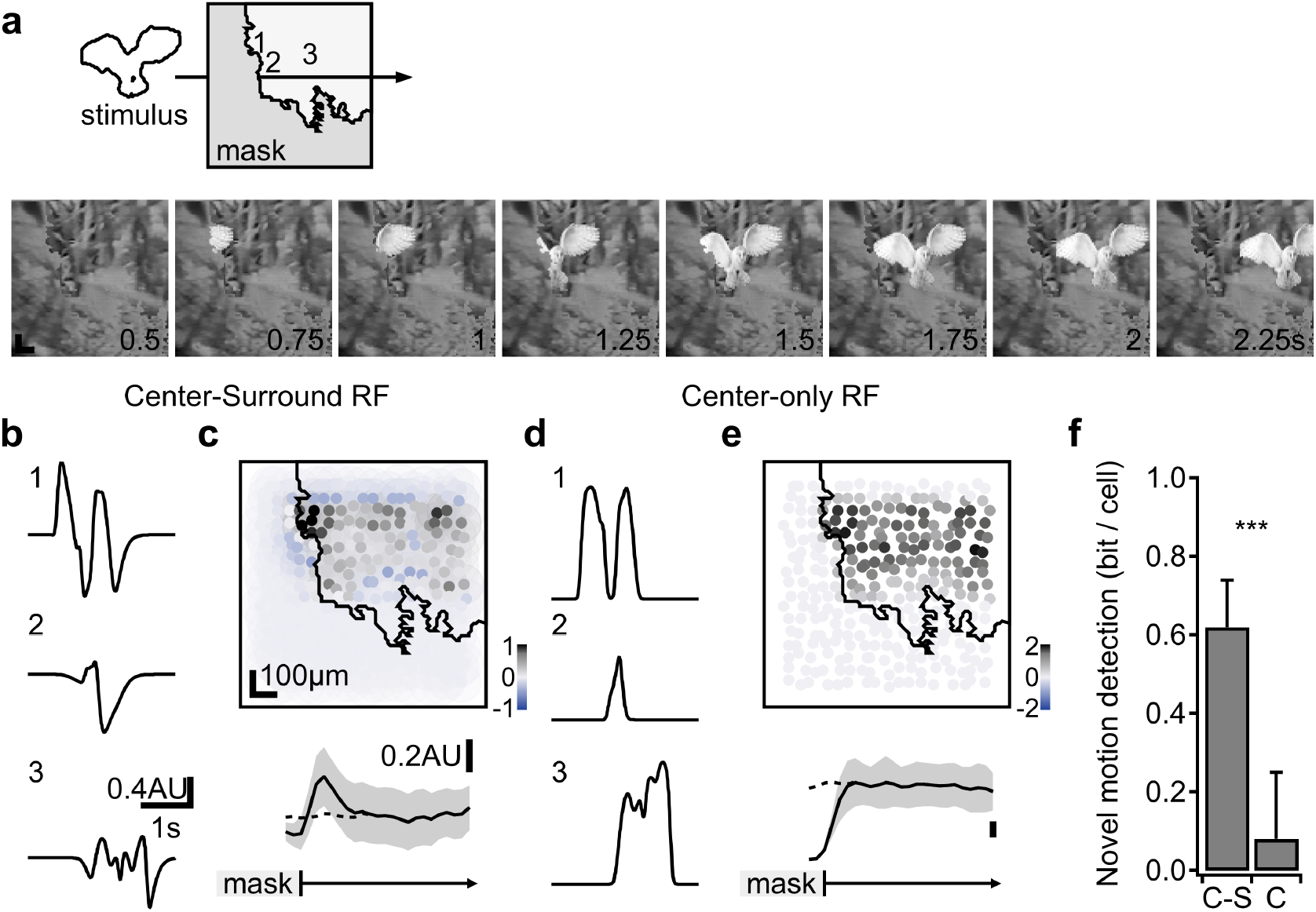
Novel object detection by linear center-surround RFs under natural movies. Simulated neuronal responses to a movie showing predator appearance (**a**). **b** Temporal response profile of three sample cells with a linear center-surround RF formulation at the spatial coordinates shown in (**a**). **c** Top, the peak response amplitude to stimulus motion from a population of simulated neurons. Activation is maximal near stimulus emergence. Bottom, the mean (±SD) change in RF activation vs. distance from the mask (n=1000 permutations of the background, the horizontal scale is preserved for both plots). Dashed trace, responses in the absence of the mask. **d-e** As in (**b-c**), with the surround component removed from the RF description. **f** The mean(±SD) mutual information computed from the differences in responses of individual neurons located near (<100 μm) the location of stimulus emergence to simulations in the presence or the absence of the mask. ***p<0.001 between the two RF architectures (t-test).

Using the simulation, we were able to test the contribution of the surround to this computation. We reformulated the RF description for the tested population to exclude the surround. We found that the outputs of the cells in this simulation were still tuned to the local contrast (Fig. 5d-e). However, the response amplitudes were similar for continuously moving and emerging stimuli, indicating that similar to our findings in the simulated retinal circuit, novel object detection required surround participation (Fig. 5e, s10).

What is the benefit of utilizing the center-surround architecture to compute novel object appearance in a realistic environment? Stronger activation near the mask-stimulus boundary can be beneficial for detecting stimuli in downstream neurons. To quantify the information that is encoded by individual neurons in our simulation, we measured the mutual information from responses of cells at the location of stimulus emergence. Analysis of signal entropies calculated from the peak responses to continuous and novel motion revealed that each cell is capable of transmitting 0.62±0.12 bits in each trial (Fig. 5f). Comparable information levels were found for responses in cells near vs. far (>200 μm) from the stimulus emergence region within the same simulation trial (data not shown). A similar analysis in center-only neurons failed to find evidence of information transfer about novel object appearance (mutual information=0.08±0.17 bits/cell, p>0.6 vs. 0, Fig. 5f), suggesting that in this scenario, postsynaptic circuits have to employ different processing schemes to detect the presence of new objects.

### Edge effects influence the analysis of motion processing in the retina

Given the participation of BCs in novel motion detection, we asked whether the dependence of BC signals on the direction of motion near mask-stimulus boundaries impacts the computation of direction selectivity (DS). The earliest direction-selective signals are present in dendrites of starburst amacrine cells (SACs), which are tuned to detect stimulus motion towards dendritic tips (Fig. 6a)^29–33^. Despite intense effort, explaining the biological implementation of this computation remains elusive^17,30,34–39^. A common strategy to probe SAC DS is to isolate dendritic computations^6,40^ with visual protocols structured to stimulate a part of the SAC^30,36,39,41^ - effectively masking part of the stimulus (Fig. 6b). To explore whether the glutamatergic drive to SACs is affected by the mask-stimulus boundary, we expressed iGluSnFR driven by the ChAT promoter (Fig. 6a)^34,35^. We presented visual stimuli as above (Fig. 6b) and set the size of the field of view to match the span of BC innervation of a single SAC dendrite (Fig. 6b, ~80 μm)^30,39^.

**Fig. 6.**
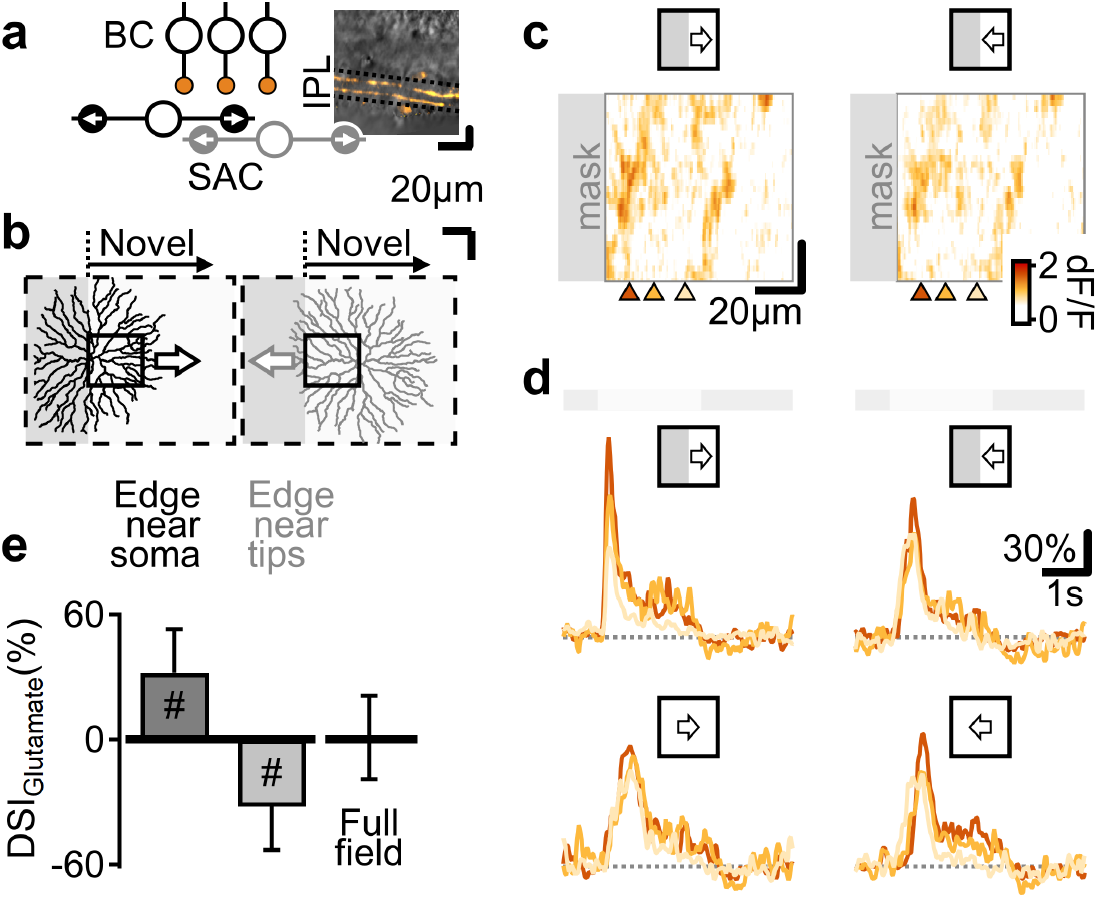
Sensitivity to novel objects in BCs influences the analysis of motion processing in postsynaptic circuits. **a** Starburst amacrine cells (SACs) integrate BC signals to detect motion towards dendritic tips. Inset, floxed-iGluSnFR expression (orange) in a Chat-Cre mouse. **b** Illustration of novel emergence enhancement vs. dendritic direction preference (arrows) in two example SACs (reconstructed in a separate experiment). Solid rectangle, the imaging window. **cd** Example peak fluorescence (**c**) and signals in ROIs (**d**, horizontal location marked in **c**) evoked by moving bars. **e** The mean (±SD) directional preference from the perspective of the SACs in **b**. #p<0.001 vs. zero.

Akin to our other findings, glutamate responses were more pronounced for emerging stimuli (Fig. 6c-d). The mean (±SD) direction selectivity index (DSI) computed from moving bar responses with the direction of motion towards/away from the boundary at the center of the display was 32 ± 21% (p<0.001 vs. 0, t-test, n=81 ROIs, Fig. 6e), while full-field moving stimuli evoked comparable glutamatergic responses in all directions (Fig. 6d-e). Could the directional effect observed in the presence of a mask-stimulus boundary contribute to DS computations in SACs? A simple analysis shows that the answer is no. The enhancement of BC drive aligns with the preferred dendritic axis in SACs whose cell bodies happen to lie near the mask (Fig. 6b, e ‘Edge near soma’). However, signals to SACs in less optimal configurations are in the ‘wrong’ direction. The grey-colored SAC illustrated in Fig. 6b serves as an example of a cell whose soma is located deeper in the stimulated region yet proximal enough to extend its dendrites over the mask (‘Edge near tips’). With the direction of motion away from the mask-stimulus boundary and towards the soma of this cell, stronger responses to emerging motion lead to a reversed directional tuning (Fig. 6e ‘Edge near tips’), in contrast to what is expected of a proper directional mechanism.

## Discussion

Using the retinal BCs as a model system, we were able to investigate the properties of motion processing in center-surround RFs. We found that the representation of continuous motion was associated with reduced peak amplitudes and prolonged temporal dynamics of glutamate signals compared with sudden object appearance in most BC types. Motion responses could not be reliably predicted from the dynamics of responses to stationary flashes, indicating a multiplexed representation of static and moving objects (Fig. 1). Visual processing in the retina is thought to be facilitated by parsing the sensory input into parallel information channels at the level of the BCs^8^. According to the literature, these communication channels represent different transformations of the photoreceptor signals and emerge from the underlying neuronal infrastructure. For example, luminance and chromatic selectivity arise from specific targeting of bipolar dendrites to distinct photoreceptors, and response polarity depends on the composition of glutamate receptors. Here, breaking with the established rules, we reveal an additional layer of complexity present in the retina. We demonstrate that individual BC types convey different temporal features of the stimulus contingent on the presence of motion. This multi-layered decomposition of the visual scene could potentially reduce the number of cells required for effective visual processing and complicates the analysis of motion responses from stationary stimuli, as is discussed in more detail below.

The unexpected diversity of motion responses revealed by our experiments highlights an asymmetric interaction between moving stimuli and static occluders; the encoding of object disappearance was similar to continuous motion, whereas newly emerging stimuli exhibited faster and more pronounced signals that qualitatively resembled the response to static flashes - particularly in transient BCs (Fig. 2).

Conceptually, our findings reflect a previously unappreciated capacity of center-surround RFs to signal the appearance of new objects. This property is a logical but previously undescribed consequence of the classic center-surround RF formulation. The mechanistic explanation for this function is straightforward and relies on the sequence of RF activation by the stimulus. Continuously moving stimuli always enter the surround RF region first, priming the surround towards a more effective inhibition by the time the center is engaged. This process doesn’t require any specific neuronal infrastructure and is present even in a linear RF formulation (Figs. 4, 5). Priming of the surround is weaker or absent in emerging motion and suddenly appearing objects. Correspondingly, the responses to these stimuli reflect the stronger role of the center component in RF integration, leading to empirically observed enhanced response amplitudes and distinct temporal dynamics between the novel and existing objects.

In our hands, the computation of novel object detection was highly prominent across all BCs, whereas their spatial contrast sensitivity, as measured by the ratio between the responses to static edges vs. full-field illumination, was not statistically significant (Fig. 2). These results suggest that in contrast to the prevailing view, sensitivity to spatial contrasts serves a secondary functional role in center-surround RFs, at least in the cells and the visual conditions we probed.

Our experiments and detailed circuit models show that photoreceptors and horizontal cells are the only circuit elements required to generate motion responses in BCs (Figs. 3, 4), leading to the conclusion that the major steps in the computation of object motion already occur at the first synapse in the retina. Our findings support the idea that signal transformation from the photoreceptors to BCs could be nonlinear and that the degree of nonlinearity is larger for transient BCs^22^. Why is processing linearity correlated with the shape of the response? Our model of signal integration in the outer plexiform layer suggests a possible answer. Nonlinear signal transformation at the photoreceptor-BC synapse could impose a threshold on the amplitude of the photoreceptor output that is required for effective activation of the postsynaptic cell (Fig. s7). As photoreceptors typically respond to light onset and light offset with a rapid membrane potential fluctuation^42,43^, nonlinear BCs are more likely to be disproportionally sensitive to these phases of photoreceptor release; their fast temporal dynamics reflect the transient shape of the filtered photoreceptor output they sample. Meanwhile, a linear signal processing mirrors the original shape of the photoreceptor light response (Figs. 4, s7). The exact biological implementation of the nonlinear photoreceptor-BC synapse dynamics is currently unclear but could plausibly be mediated by a differential affinity of BC dendrites to photoreceptor release^44^. In the end, the nonlinear nature of the transient BC population is known to contribute to a rudimentary feature detector-like behavior that is tuned to certain visual conditions, such as signal polarity and spatial inhomogeneity^22^. We can now add novel object appearance to this list.

We used stimuli that were explicitly designed to compare responses to moving and static objects and representation of novel vs. existing visual items. Previous reports demonstrated that the retina is capable of detecting acceleration^5^, differential motion^45^, looming (approaching) motion^4^, and distinguish between a motion to uncorrelated spatiotemporal activation^3^. These visual functions are thought to require higher-order retinal neurons (e.i., amacrine and ganglion cells). How RF components are integrated during the presentation of these stimuli and whether RF computations contribute to such sophisticated calculations remain to be elucidated.

In the last decade, several groups found evidence for a spatial offset between presynaptic BC populations that are aligned with the directional axis in DS ganglion cells and dendritic position in SACs^17,24,30,46^. This circuit organization can support directional tuning by a mechanism first described by Hassenstein and Reichardt^47^ - if the response speed of the BCs follows their spatial arrangement. Conflicting results were reached in studies designed to test the predictions of this model using electrophysiological and imaging approaches^18,39,48^. Importantly, all previous work examined BC output in response to the presentation of stationary inputs, which, as our results indicate, do not accurately reflect the dynamics in BCs during motion. Proposed directional computations are particularly dependent on BCs rise times, which, as our data reveal, are uncorrelated between moving and static objects (Fig. 1). At the very least, the dramatic increase in the rise-times dynamics we observed in our recordings suggests a shift in speed dependence of the Hassenstein-Reichardt detector to slow-moving objects. Further experiments will be required to resolve this issue and elucidate the potential impact of visual edges on observed DS (Fig. 6).

Our findings of motion processing in the early processing stages in the retina have intriguing psychophysical implications to the perception of novel stimuli over continuing motion and echo the salience of visual perception in humans^49,50^: the sudden appearance of new objects grabs attention reflexively; motion onset is less salient - but more noticeable than continuous motion. Our data propose that these computations are hard-wired in the retina and reflect the information content conveyed by the respective visual items. From an ecological perspective, the utility of continuous retinal motion is diminished as it may be self-generated by locomotion through the environment and because the trajectory of continuously moving objects could be predicted by past sensory input. Conversely, novel stimuli can alert to a predator or prey; their fast processing is vital to survival. All the necessary machinery for motion processing in BCs we describe in the mouse are conserved in primates, providing strong evidence that enhanced representation of newly flashed and emerging moving objects are consequences of a bottom-up process fundamental to how visual stimuli are computed in the retina.

Taken together, our work complements previous studies revealing decorrelation of signals by surround inhibition^9,21,51^ and shows how simple operational concepts give rise to complex visual computations. Diverse representation of different features of the visual space in a single neuronal population and early detection of salient environmental cues are powerful strategies that reduce the computational burden of the visual system. The surprising finding that the classic center-surround RF architecture is sufficiently versatile to take part in seemingly unrelated tasks is critical to the understanding of visual computations in multiple brain regions and the design of future studies of visual perception.

## Methods

### Virus expression

All animal procedures were conducted in accordance with U.S. National Institutes of Health guidelines, as approved by the University of Colorado Institutional Animal Care and Use Committee (IACUC). For intravitreal virus injections, mice of ages p21–120 of either sex were anaesthetized with isoflurane; ophthalmic proparacaine and phenylephrine were applied for pupil dilation and analgesia. A small incision at the border between the sclera and the cornea was made with a 30 gauge needle. 1 μL of AAV solution was injected with a blunt tip (30 gauge) modified Hamilton syringe (http://retina.anatomy.upenn.edu/~bart/B_Sci/InjectorSyringepayments.html). AAV9.hsyn.iGluSnFR. WPRE.SV40, (a gift from Loren Looger, Addgene plasmid # 98929; http://n2t.net/addgene:98929; RRID:Addgene_98929; 10^13^ vg/mL in water) was injected into the vitreous humour of wild type mice (C57BL/6J, Jackson laboratory, www.jax.org). To express iGluSnFR in SACs only, AAV9.hsyn.FLEX.iGluSnFR.WPRE.SV40 (a gift from Loren Looger, Addgene plasmid # 98931; http://n2t.net/addgene: 98931; RRID:Addgene_98931, similar concentration) was used in Chat-Cre transgenic mice. AAV9-pGP-AAV-syn-jGCaMP7f-WPRE (Addgene plasmid # 104488; http://n2t.net/addgene: 104488; RRID:Addgene_104488) was used to measure intracellular calcium levels. Experiments on retinas from all animal groups were performed 2-6 weeks following virus injection.

### Imaging procedures

Mice were not dark-adapted to reduce rod-pathway activation. Two hours after enucleation, retina sections were whole mounted on a platinum harp with their photoreceptors facing down, suspended ~1 mm above the glass bottom of the recording chamber. The retina was kept ~32°C and continuously superfused with Ames media (Sigma-Aldrich, www.sigmaaldrich.com) equilibrated with 95%O_2_/5%CO_2_.

### Light-stimulation

Light stimuli were generated in Igor Pro 8 (Wavemetrics, www.wavemetrics.com) PC and displayed with a 415 nM LED collimated and masked by an LCD display (3.5 Inch, 480×320 pixels, refresh rate of 50 Hz) controlled by a custom-written python script running on raspberry pi 3 computer. Display luminosity was gamma corrected with a powermeter (Thorlabs, www.thorlabs.com); the stimulus was set to either 60% or −60% Michelson contrast. Frame timing was controlled by a clock signal from Sutter IPA patch-clamp amplifier (Sutter Instruments, www.sutter.com) driven by Igor Pro and read from one of the digital I/O ports of the raspberry pi. Light from the visual stimulus was focused by the condenser to illuminate the tissue at the focal plane of the photoreceptors (resolution = 2.5 μm/pixel, background light intensity = 30,000-60,000 R* rod^-1^). Both vertical and horizontal light stimulus positions were checked and centered daily before the start of the experiments. The following light stimulus patterns were used: static bar covering the entire display (800×800 μm) presented for 2 seconds. A 1 mm-long bar moving either to the left or the right directions (speed = 0.5 mm/s; dwell time over each pixel = 2 s). These stimuli were repeated with masks (at background light levels), spanning the full height of the display, occluding different portions of the stimulus. Each visual stimulation protocol was repeated at least 3 times.

### Imaging

Glutamate and calcium imaging was performed with Throlabs Bergamo galvo-galvo two-photon microscope. A pulsed laser light (920 nm, ~1 μW output at the objective; Chameleon Ultra II, Coherent, www.coherent.com) was used for two-photon excitation projected from an Olympus 20X (1 NA) objective. A descanned (confocal) photomultiplier tube (PMT) was used to acquire fluorescence between 500 and 550 nm. The confocal pinhole (diameter = 1 mm) largely prevented stimulus light (focused on a different focal plane), from reaching the PMT, allowing us to present the visual stimulus during two-photon imaging. A photodiode mounted under the condenser sampled transmitted laser light to generate a reference image of the tissue. Fluorescence signals were collected in a rapid bidirectional frame scan mode (128×64 pixels; ~50 Hz, Thorimage). The line spacing on the vertical axis was doubled to produce a rectangular imaging window (typically ~82×82 μm size, in some experiments, the window was set to ~164×164 μm; the corresponding pixel sizes were 0.64 μm or 1.28 μm). To reduce shot noise, images were subsampled by averaging 2×2 neighboring pixels and filtered by a 20 Hz low pass filter offline. Horizontal and vertical image drifts were corrected online using a reference z-stack acquired before time-series recordings.

For pharmacological manipulations, we used SR95531 (50 μM, Abcam, www.abcam.com) to block GABA_A_ receptors, TPMPA (50 μM, Tocris, www.tocris.com) to block GABAC receptors and strychnine (1 μM), Abcam) to block glycine receptors. All drugs were mixed with the bath Ames medium.

### Analysis

All analysis was done in Igor Pro 8. Fluorescence signals were averaged across repeated visual protocol presentations. Pixels with dF/F values >20% were selected for clustering analysis. For the initial clustering of ROIs with similar response kinetics, we combined 1-second recordings of the response shapes around the time of stimulus entrance to the imaging window from each of the tested visual protocols across all imaged planes. A similarity matrix was constructed from a pairwise pixel comparison measured with Igor build-in farthest-point clustering algorithm. McClain-Rao index was used to determine the optimal number of clusters^32^. The shapes of the resulting ROIs were fitted with a sigmoid for the rising phase of the response and with a single exponential for the decay phase. ROIs were manually curated and removed from analysis if pixel variability, measured with a coefficient of variation, exceeded 1.

We computed the horizontal RF position from responses to motion over the entire display. We first determined the timing of 50% rise-time from trials with leftward and rightward motion. ROIs with their RF center in the middle of the display should respond to both stimuli at the same time following stimulus presentation. In an ROI where the center of the RF is located to the left/right of the display center, a rightward moving stimulus elicits a response that comes earlier/later compared to a trial with a leftward moving stimulus. RF position was computed as half the time difference between the diametrically opposed trials, multiplied by stimulus speed. Trial responses were considered to be to full-field stimulation if the RF center was at least 100 μm away from the nearest visual edge formed either by masks or the boundaries of the display. Similarly, responses were considered to be near an edge if at least one of the visual edges was closer than 50 μm to the RF center.

To detect similarly shaped groups between different experiments, we conducted a secondary hierarchical clustering. Our initial clustering incorporated responses from trials with moving stimuli and responses near visual edges. Motion responses shift in time as the stimulus progresses over the retina, making comparisons between ROIs difficult. Edge effects may also affect the shape of the responses. For these reasons, as an input to the similarity matrix, we performed a pairwise comparison between 1-second long responses to full-field static stimulation only, for positive contrast stimuli presentation for ON groups and negative stimuli for the OFF groups. As before, the optimal cluster number was determined with the McClain-Rao index analysis.

Transiency index (TI) was calculated as the ratio between the peak and the mean of the response within the stimulation window. TI=1 indicates a sharp and transient response, TI close to zero is produced by sustained plateaus.

Direction Selectivity Index (DSI) was calculated as a vector sum of vectors V_i_ pointing in the direction of the stimulus and having the length R_i_ = peak dF/F of the response to that stimulus.

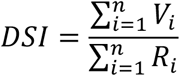

Where n is the number of probed directions. DSI can range from 0 to 1, with zero indicating no directional preference and 1 indicating responses to only one direction of stimulation.

Bonferroni correction was used for multiple comparisons. Whenever ratios between parameters were compared, statistics were computed on a logarithmic transformation of the data.

### Modeling

All simulations were conducted in Igor Pro 8.

### Linear receptive field model

We simulated a simple spatiotemporal RF structure to examine the engagement of a cell with a center-surround RF organization by visual motion. The spatial extent of the center and surround RF components were defined by a two-dimensional Gaussian function with half widths of 50 and 200 μm, respectively. The responses for the RF components were modeled as a single exponential with a time constant (τ) of 20 ms for the center and 100 ms for the surround. The simulation ran for 4500 ms with a time step of 1 ms. In each step, the total illuminated RF area was computed from the convolution of the center/surround RF components with the stimulus. RF activation at time step *t* was changed by the difference between the sum of the newly illuminated RF area and the signal from the previous time step:

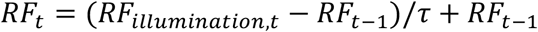

The full RF was computed according to the following equation:

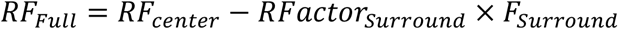

Where Factor_surround_ indicated the intensity of the surround activation and varied between 0 (no surround) to 0.5.

Simulated neurons were distributed on a 1000 x 1000 μm square grid stimulated either by moving / stationary bars with similar parameters (speed, contrast, size) as in the experiments or by natural images.

### Natural movies

The natural movies were composed of background/mask chosen from individual frames of the ‘catcam’ database^22,52^ and stimuli depicting birds of prey. The images were cropped to 100 x 100 pixels and presented as an input to the simulated network. The intensity of the background/mask was scaled to be at the mean pixel level (i.e., 128 pixel luminance value) with an SD of 30. The mean intensity of the stimuli was set to be 2 SD higher than the background mean. In some simulations, the stimulus was not presented. Instead, the background translated horizontally at 0.5 mm/s as measured over the artificial retina. The shape of the mask was chosen by foreground objects in separate movie frames. The mask was absent for simulations of continuous motion. Response amplitudes were measured in a time window spanning 500 ms starting at the time of object appearance over the location of the simulated cell. Mutual information was measured as the entropy of responses near (<100 μm) the initial appearance of the stimulus near the mask/stimulus boundary in the presence/absence of the mask, minus the average entropy of responses to the individual conditions.

### Detailed retinal simulation

The simulated retina consisted of a one-dimensional array (length=700 μm) of photoreceptors, horizontal cells, and BCs, spaced 10 μm apart. Stimuli were provided by a bright bar that was either flashed for 2 seconds or moved over the retina (speed=0.5 mm/s). Visual edges were created by masking visual presentation near the borders of the array. The simulation time step was 1 ms.

Photoreceptor activation was modeled as a difference between two activation functions (Ph_A_, Ph_B_) with instantaneous rise time and decay times of 60 and 400 ms, respectively.

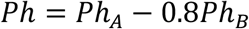

Time step computations for the activation functions were given by:

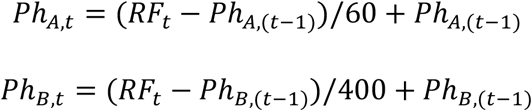

Where RF was computed from the value of the stimulus at the position of the photoreceptor and horizontal cell feedback (see below) and Pht-1 represents the value of the activation function on a previous time step.

Horizontal cells integrated all photoreceptor signals in their RF. The spatial RF signal in horizontal cell_*i*_ (HC_∞,i_) was described by a projection of a two-dimensional Gaussian function with a radius of 60 μm on the single spatial dimension of the photoreceptor array according to the following:

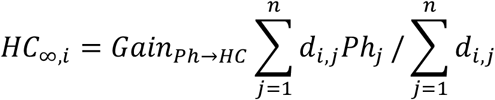

Where the Photoreceptor-horizontal cell gain was set to 1 unless specified otherwise; n=150 is the number of photoreceptors, d_*i,j*_Ph_*j*_ represents the dimensionality corrected signal from photoreceptor *j* on horizontal cell_*i*_ and the last term used to correct responses by RF size.

The total activation of the horizontal cells at a time step *t* was given by the following equation:

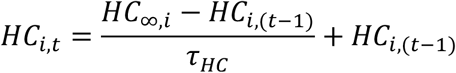

In which τ_HC_ is the horizontal cell activation time constant = 120 ms.

Each photoreceptor combined horizontal cell signals (normalized by the same distance) function) with visual illumination as follows:

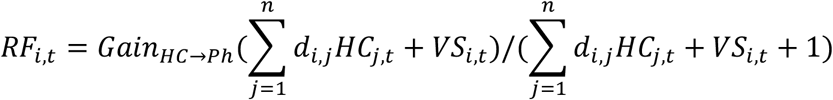

Where the photoreceptor-horizontal cell gain was set to 1, VS_*i,t*_ represents the value of the visual stimulus over photoreceptor *i* at time t and HC_*j,t*_ is the feedback from horizontal cell *j*.

Similar to horizontal cells, BCs sampled photoreceptor input by dimensionality-corrected RF (size=50 μm unless specified otherwise). The steady-state input-output transformation at the photoreceptor-BC synapse was given by the following relationship:

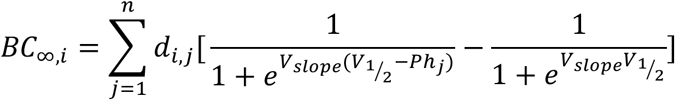

Where d_*i,j*_ was the distance function computed as for horizontal cells, V_slope_ and V_½_ defined the slope and the 50% point of the Ph-BC transformation function, and the last term provided a subtraction of the baseline photoreceptive signal.

Last, the actual voltage at each BC *i* at time step *t* was computed using the following:

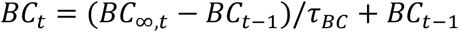

In which τ_BC_ indicate the activation time constant = 60 ms (unless specified otherwise).

## Author contributions

**John Gaynes:** Investigation, Writing-Review &Editing, Supervision **Samuel Budoff:** Investigation, Formal analysis, Writing-Review &Editing **Michael Grybko:** Investigation, Writing-Review &Editing **Joshua Hunt:** Formal analysis, Writing-Review &Editing **Alon Poleg-Polsky:** Conceptualization, Methodology, Software, Formal analysis, Resources, Data Curation, Writing-Original draft, Supervision, Project administration, and Funding acquisition

## Data and materials availability

The code for the visual stimulation and simulations is available on (https://github.com/PolegPolskyLab/). The full dataset has not been uploaded in a public repository due to file size limitation but is available upon request from.

## References

1 Zhang, Y., Kim, I. J., Sanes, J. R. & Meister, M. The most numerous ganglion cell type of the mouse retina is a selective feature detector. Proceedings of the National Academy of Sciences of the United States of America 109, E2391–2398, doi:10.1073/pnas.1211547109 (2012).

2 Olveczky, B. P., Baccus, S. A. & Meister, M. Segregation of object and background motion in the retina. Nature 423, 401–408, doi:10.1038/nature01652 (2003).

3 Manookin, M. B., Patterson, S. S. & Linehan, C. M. Neural Mechanisms Mediating Motion Sensitivity in Parasol Ganglion Cells of the Primate Retina. Neuron, doi:10.1016/j.neuron.2018.02.006 (2018).

4 Munch, T. A. et al. Approach sensitivity in the retina processed by a multifunctional neural circuit. Nature neuroscience 12, 1308–1316, doi:10.1038/nn.2389 (2009).

5 Schwartz, G., Taylor, S., Fisher, C., Harris, R. & Berry, M. J., 2nd. Synchronized firing among retinal ganglion cells signals motion reversal. Neuron 55, 958–969, doi:10.1016/j.neuron.2007.07.042 (2007).

6 Mauss, A. S., Vlasits, A., Borst, A. & Feller, M. Visual Circuits for Direction Selectivity. Annu Rev Neurosci, doi:10.1146/annurev-neuro-072116-031335 (2017).

7 Wei, W. Neural Mechanisms of Motion Processing in the Mammalian Retina. Annu Rev Vis Sci 4, 165–192, doi:10.1146/annurev-vision-091517-034048 (2018).

8 Euler, T., Haverkamp, S., Schubert, T. & Baden, T. Retinal bipolar cells: elementary building blocks of vision. Nature reviews 15, 507–519, doi:10.1038/nrn3783 (2014).

9 Franke, K. et al. Inhibition decorrelates visual feature representations in the inner retina. Nature 542, 439–444, doi:10.1038/nature21394 (2017).

10 Behrens, C., Schubert, T., Haverkamp, S., Euler, T. & Berens, P. Connectivity map of bipolar cells and photoreceptors in the mouse retina. eLife 5, doi:10.7554/eLife.20041 (2016).

11 Baden, T., Berens, P., Bethge, M. & Euler, T. Spikes in mammalian bipolar cells support temporal layering of the inner retina. Current biology: CB 23, 48–52, doi:10.1016/j.cub.2012.11.006 (2013).

12 Baden, T., Euler, T. & Berens, P. Understanding the retinal basis of vision across species. Nature reviews, doi:10.1038/s41583-019-0242-1 (2019).

13 Borghuis, B. G., Marvin, J. S., Looger, L. L. & Demb, J. B. Two-photon imaging of nonlinear glutamate release dynamics at bipolar cell synapses in the mouse retina. The Journal of neuroscience: the official journal of the Society for Neuroscience 33, 10972–10985, doi:10.1523/JNEUROSCI.1241-13.2013 (2013).

14 Baccus, S. A., Olveczky, B. P., Manu, M. & Meister, M. A retinal circuit that computes object motion. J Neurosci 28, 6807–6817 (2008).

15 Schwartz, G. W. et al. The spatial structure of a nonlinear receptive field. Nature neuroscience 15, 1572–1580, doi:10.1038/nn.3225 (2012).

16 Thoreson, W. B. & Mangel, S. C. Lateral interactions in the outer retina. Progress in retinal and eye research 31, 407–441, doi:10.1016/j.preteyeres.2012.04.003 (2012).

17 Kim, J. S. et al. Space-time wiring specificity supports direction selectivity in the retina. Nature 509, 331–336, doi:10.1038/nature13240 (2014).

18 Fransen, J. W. & Borghuis, B. G. Temporally Diverse Excitation Generates Direction-Selective Responses in ON- and OFF-Type Retinal Starburst Amacrine Cells. Cell reports 18, 1356–1365, doi:10.1016/j.celrep.2017.01.026 (2017).

19 Turner, M. H., Schwartz, G. W. & Rieke, F. Receptive field center-surround interactions mediate context-dependent spatial contrast encoding in the retina. eLife 7, doi:10.7554/eLife.38841 (2018).

20 Kuo, S. P., Schwartz, G. W. & Rieke, F. Nonlinear Spatiotemporal Integration by Electrical and Chemical Synapses in the Retina. Neuron 90, 320–332, doi:10.1016/j.neuron.2016.03.012 (2016).

21 Pitkow, X. & Meister, M. Decorrelation and efficient coding by retinal ganglion cells. Nature neuroscience 15, 628–635, doi:10.1038/nn.3064 (2012).

22 Schreyer, H. M. & Gollisch, T. Nonlinear spatial integration in retinal bipolar cells shapes the encoding of artificial and natural stimuli. Neuron, doi:10.1016/j.neuron.2021.03.015 (2021).

23 Marvin, J. S. et al. An optimized fluorescent probe for visualizing glutamate neurotransmission. Nat Methods 10, 162–170, doi:10.1038/nmeth.2333 (2013).

24 Matsumoto, A., Briggman, K. L. & Yonehara, K. Spatiotemporally Asymmetric Excitation Supports Mammalian Retinal Motion Sensitivity. Curr Biol, doi:10.1016/j.cub.2019.08.048 (2019).

25 Hartline, H. K., Wagner, H. G. & Ratliff, F. Inhibition in the eye of Limulus. The Journal of general physiology 39, 651–673, doi:10.1085/jgp.39.5.651 (1956).

26 Marr, D. & Hildreth, E. Theory of edge detection. Proceedings of the Royal Society of London. Series B, Containing papers of a Biological character 207, 187–217, doi:10.1098/rspb.1980.0020 (1980).

27 Grove, J. C. R. et al. Novel hybrid action of GABA mediates inhibitory feedback in the mammalian retina. PLoS biology 17, e3000200, doi:10.1371/journal.pbio.3000200 (2019).

28 Barnes, S., Grove, J. C. R., McHugh, C. F., Hirano, A. A. & Brecha, N. C. Horizontal Cell Feedback to Cone Photoreceptors in Mammalian Retina: Novel Insights From the GABA-pH Hybrid Model. Front Cell Neurosci 14, 595064, doi:10.3389/fncel.2020.595064 (2020).

29 Euler, T., Detwiler, P. B. & Denk, W. Directionally selective calcium signals in dendrites of starburst amacrine cells. Nature 418, 845–852 (2002).

30 Ding, H., Smith, R. G., Poleg-Polsky, A., Diamond, J. S. & Briggman, K. L. Species-specific wiring for direction selectivity in the mammalian retina. Nature 535, 105–110, doi:10.1038/nature18609 (2016).

31 Morrie, R. D. & Feller, M. B. A Dense Starburst Plexus Is Critical for Generating Direction Selectivity. Curr Biol 28, 1204–1212 e1205, doi:10.1016/j.cub.2018.03.001 (2018).

32 Poleg-Polsky, A., Ding, H. & Diamond, J. S. Functional Compartmentalization within Starburst Amacrine Cell Dendrites in the Retina. Cell reports 22, 2898–2908, doi:10.1016/j.celrep.2018.02.064 (2018).

33 Yonehara, K. et al. The first stage of cardinal direction selectivity is localized to the dendrites of retinal ganglion cells. Neuron 79, 1078–1085, doi:10.1016/j.neuron.2013.08.005 (2013).

34 Gavrikov, K. E., Nilson, J. E., Dmitriev, A. V., Zucker, C. L. & Mangel, S. C. Dendritic compartmentalization of chloride cotransporters underlies directional responses of starburst amacrine cells in retina. Proceedings of the National Academy of Sciences of the United States of America 103, 18793–18798, doi:0604551103 [pii] 10.1073/pnas.0604551103 (2006).

35 Hausselt, S. E., Euler, T., Detwiler, P. B. & Denk, W. A dendrite-autonomous mechanism for direction selectivity in retinal starburst amacrine cells. PLoS biology 5, e185 (2007).

36 Oesch, N. W. & Taylor, W. R. Tetrodotoxin-resistant sodium channels contribute to directional responses in starburst amacrine cells. PLoS One 5, e12447, doi:10.1371/journal.pone.0012447 (2010).

37 Koren, D., Grove, J. C. R. & Wei, W. Cross-compartmental Modulation of Dendritic Signals for Retinal Direction Selectivity. Neuron 95, 914–927 e914, doi:10.1016/j.neuron.2017.07.020 (2017).

38 Chen, Q., Pei, Z., Koren, D. & Wei, W. Stimulus-dependent recruitment of lateral inhibition underlies retinal direction selectivity. eLife 5, doi:10.7554/eLife.21053 (2016).

39 Vlasits, A. L. et al. A Role for Synaptic Input Distribution in a Dendritic Computation of Motion Direction in the Retina. Neuron 89, 1317–1330, doi:10.1016/j.neuron.2016.02.020 (2016).

40 Demb, J. B. Cellular mechanisms for direction selectivity in the retina. Neuron 55, 179–186 (2007).

41 Lee, S. & Zhou, Z. J. The synaptic mechanism of direction selectivity in distal processes of starburst amacrine cells. Neuron 51, 787–799 (2006).

42 Dunn, F. A., Lankheet, M. J. & Rieke, F. Light adaptation in cone vision involves switching between receptor and post-receptor sites. Nature 449, 603–606 (2007).

43 Clark, D. A., Benichou, R., Meister, M. & Azeredo da Silveira, R. Dynamical adaptation in photoreceptors. PLoS Comput Biol 9, e1003289, doi:10.1371/journal.pcbi.1003289 (2013).

44 DeVries, S. H., Li, W. & Saszik, S. Parallel processing in two transmitter microenvironments at the cone photoreceptor synapse. Neuron 50, 735–748, doi:10.1016/j.neuron.2006.04.034 (2006).

45 Olveczky, B. P., Baccus, S. A. & Meister, M. Retinal adaptation to object motion. Neuron 56, 689–700 (2007).

46 Greene, M. J., Kim, J. S., Seung, H. S. & EyeWirers. Analogous Convergence of Sustained and Transient Inputs in Parallel On and Off Pathways for Retinal Motion Computation. Cell reports 14, 1892–1900, doi:10.1016/j.celrep.2016.02.001 (2016).

47 Hassenstein, B., Reichardt, W. Systemtheoretische Analyse Der Zeit, Reihenfolgen Und Vorzeichenauswertung Bei Der Bewegungsperzeption Des Rüsselkäfers Chlorophanus. Z. Naturforsch. B 11, 513–524 (1956).

48 Stincic, T., Smith, R. G. & Taylor, W. R. Time course of EPSCs in ON-type starburst amacrine cells is independent of dendritic location. The Journal of physiology 594, 5685–5694, doi:10.1113/JP272384 (2016).

49 Abrams, R. A. & Christ, S. E. Motion onset captures attention. Psychol Sci 14, 427–432, doi:10.1111/1467-9280.01458 (2003).

50 Smith, K. C. & Abrams, R. A. Motion onset really does capture attention. Atten Percept Psychophys 80, 1775–1784, doi:10.3758/s13414-018-1548-1 (2018).

51 Vinje, W. E. & Gallant, J. L. Sparse coding and decorrelation in primary visual cortex during natural vision. Science (New York, N.Y 287, 1273–1276, doi:10.1126/science.287.5456.1273 (2000).

52 Betsch, B. Y., Einhauser, W., Kording, K. P. & Konig, P. The world from a cat’s perspectivestatistics of natural videos. Biological cybernetics 90, 41–50, doi:10.1007/s00422-003-0434-6 (2004).

