## supplemental files for "Novel Object Detection and Multiplexed Motion Representation in Retinal Bipolar Cells"

**Supplementary figures**

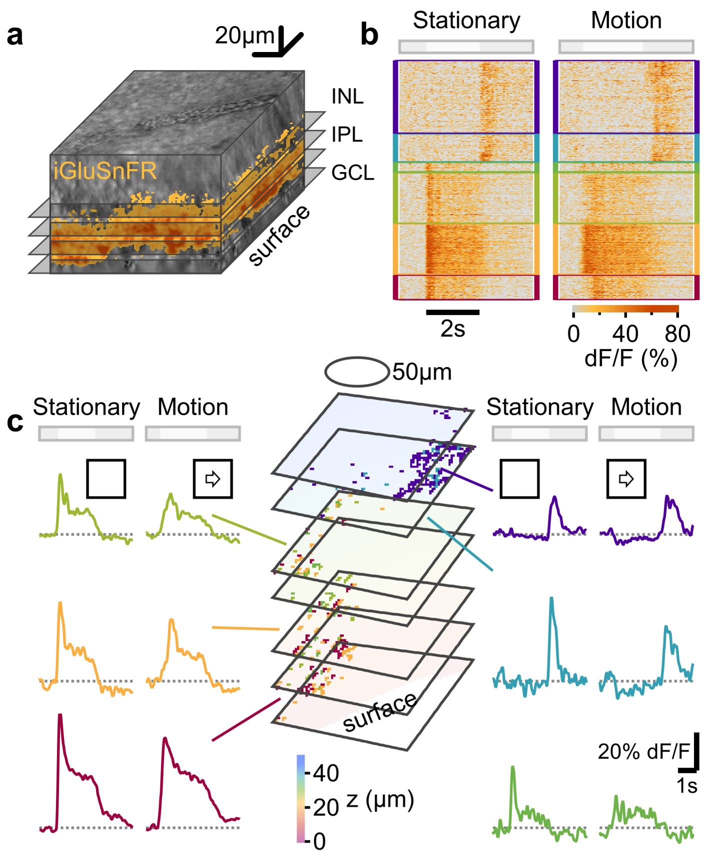


**Fig. S1. Recording and analysis of light-driven glutamatergic responses in the inner retina. a** 3D reconstruction of a two-photon imaged retina, showing iGluSnFR expression in the inner retina (orange) and a schematic of the scan planes. INL, inner nuclear layer; IPL, inner plexiform layer; GCL, ganglion cell layer. **b** A heatmap illustrating the temporal responses to full-field stationary (left) and moving (right) bars recorded from representative pixels in the retina shown in **a**. Responses were clustered into ROIs based on the similarity of response waveforms. **c** The mean signals from the ROIs outlined in **b**. The middle panel shows the location of the pixels belonging to the ROI across scan planes. Ellipse-the approximate extent of a center RF region in BCs. Color-coding based on the mean IPL depth.

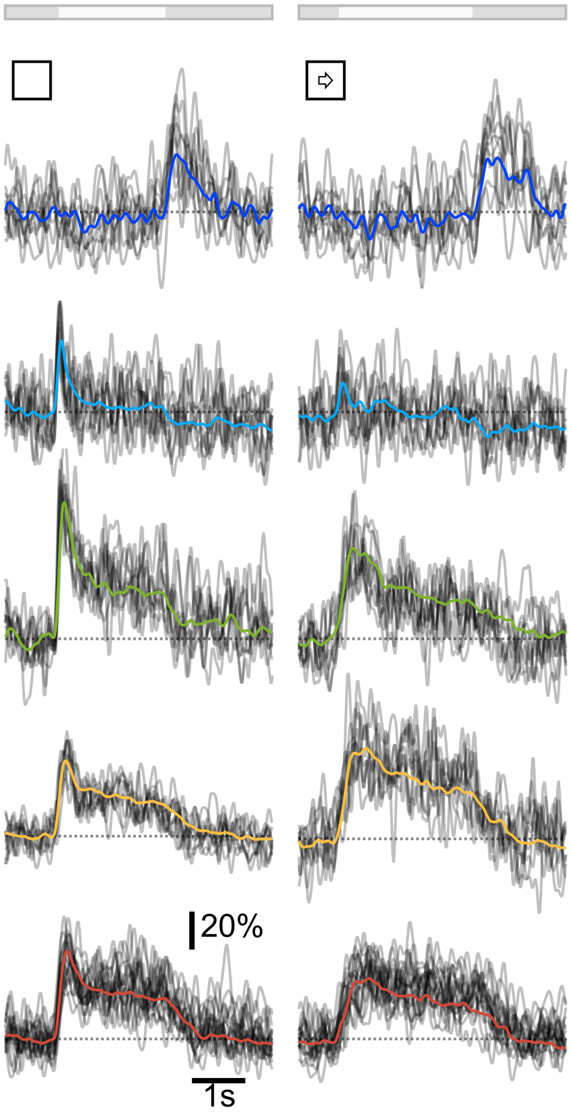


**Fig. S2. Grouping of iGluSnFR signals**

Grey, ten representative traces of fluorescence change over time, computed in individual pixels belonging to 5 different iGluSnFR ROIs. The clustering algorithm compared the similarity between responses from individual pixels in the image that passed the inclusion criteria (methods).

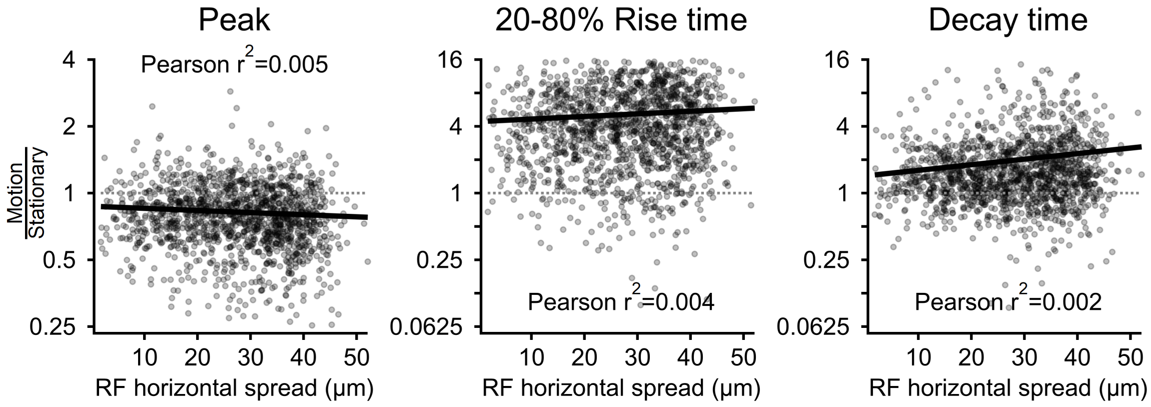


**Fig. S3. Little effect of the spatial pixels spread on the extrapolated ROI dynamics**

The ratio between the peak (left), rise time (middle) and decay time (right) responses to full-field motion vs. stationary stimulation as a function of the SD of the horizontal pixel spread for all ROIs in the dataset. Solid lines, linear fits to the data.

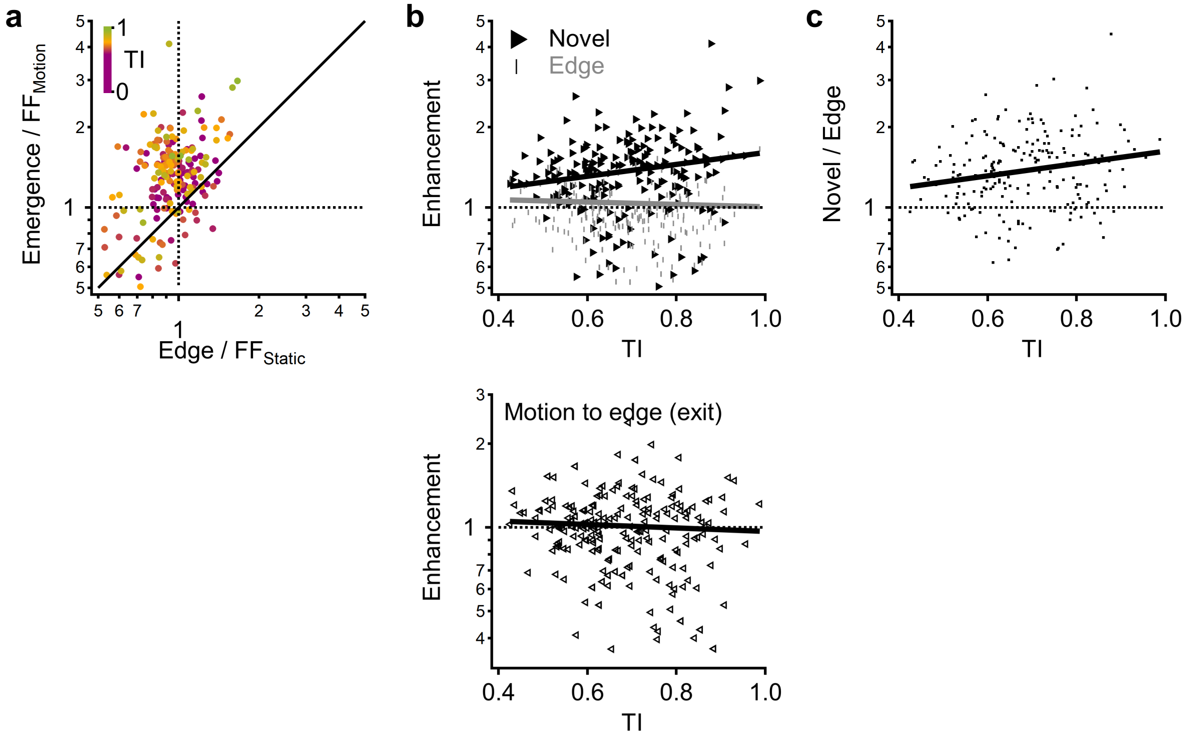


**Fig. S4. Novel motion detection is more prominent compared to edge enhancement. a** The relationship between novel object enhancement (measured as the ratio between the peak of the ROI signal to emerging object vs. full-field motion) and edge enhancement (measured as the ratio between the peak of the stationary response near mask-stimulus boundary vs. response to full-field flash). Color coding is by the transiency index calculated from the full-field static signals. **b** Top, novel object (black) and edge (grey) enhancement as a function of ROI transiency. Bottom, same for object exit (calculated as the peak ROI signal to motion towards the edge normalized by the peak of the full-field motion response). **c** The ratio between novel object and edge enhancement vs. the transiency index. Solid lines in **b** and **c** indicate linear fits. More pronounced emerging motion enhancement is observed in transient ROIs. The majority of ROIs were tuned to the detection of novel objects as compared to the presence of static edges.

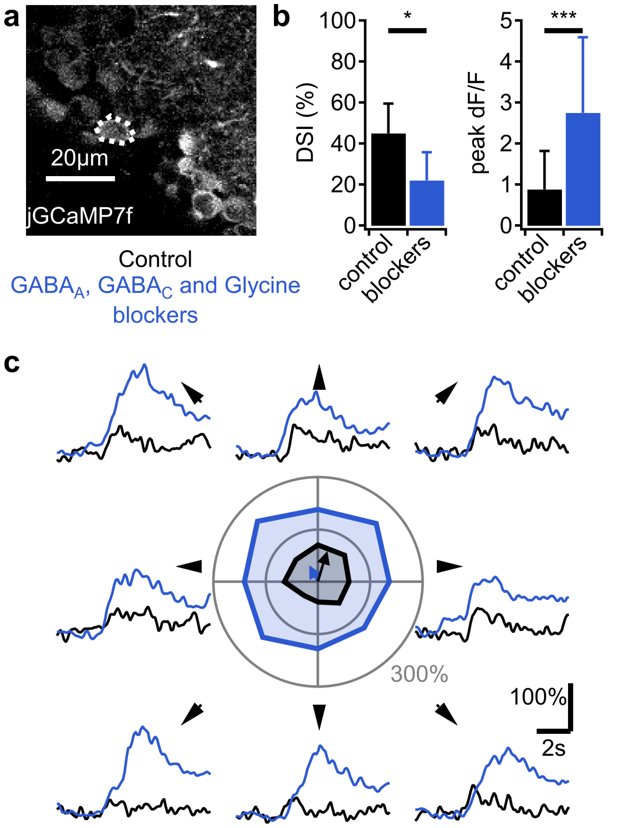


**Fig. S5. Validation of inhibitory blockers effectiveness on responses of ganglion cells**

**a** jGCAMP7f was expressed in the ganglion cell layer of WT mice and ganglion cell activity was monitored in ex-vivo retina during stimulation with bright bars moving in 8 directions separated by 45˚. Direction selective cells were identified by having a DSI>10%. Following control recording in Ames solution, blockers of $GABA_A$, $GABA_C$ and glycine were added to the bath. **b** Comparison between the DSI values of DS ganglion cells (left, n=6) and peak fluorescence in all recorded ganglion cells (right, n=22) in control solution (black) and after addition of inhibitory blockers (grey). * p<0.05,***p<0.001; paired t-test. Error bars-SD. **c** Example light responses from a DS ganglion cell outlined in **a**. The directional vector was computed as the vector sum of the peak responses to different directions. Color coding as in **b**. As expected from the role of inhibition in governing the size of DS in ganglion cells and in dampening neuronal responses in general, we observed reduced DS and larger calcium transients following the application of the drugs.

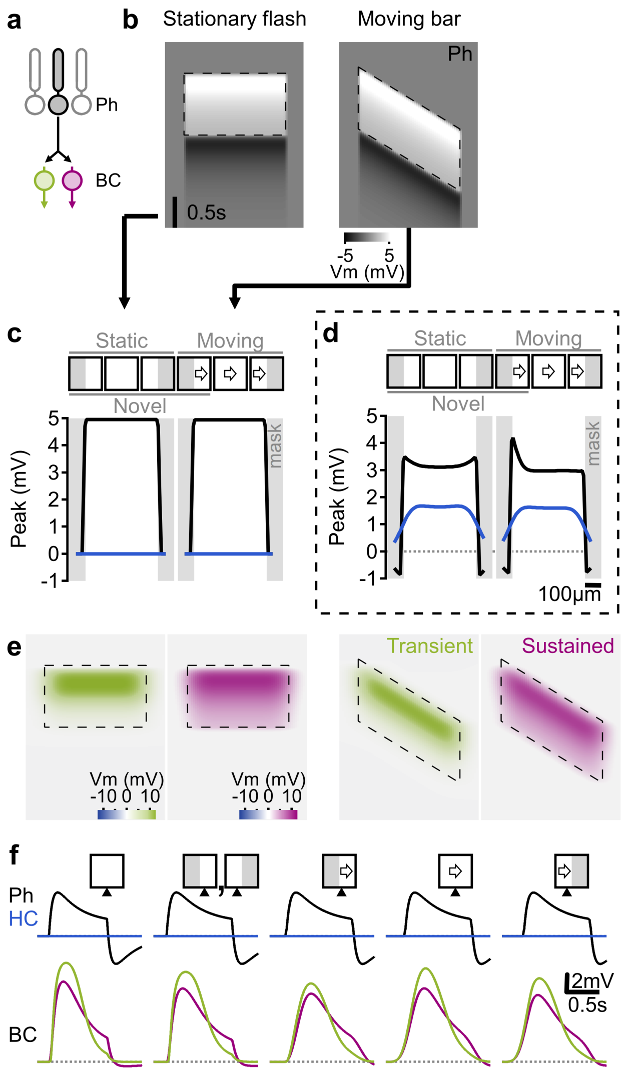


**Fig. S6. Lateral inhibition is required to establish differential responses to novel objects and edges. a** The simulated retinal circuit that did not include inhibitory components. Other network parameters and stimuli were left unchanged. **b** Space-time plot of photoreceptor activation. **c** Peak depolarization of photoreceptors (black) and HC (blue) vs. spatial positions. **d** As in **c** for the full model shown in **Fig. 4**. **e** as in **b** for transient (green) and sustained (red) BCs. **f** Example responses from cells near edges and the center of the stimulated region. The surround was not absolutely required for generating smaller and slower responses to motion. Edge effects, however, were absent.

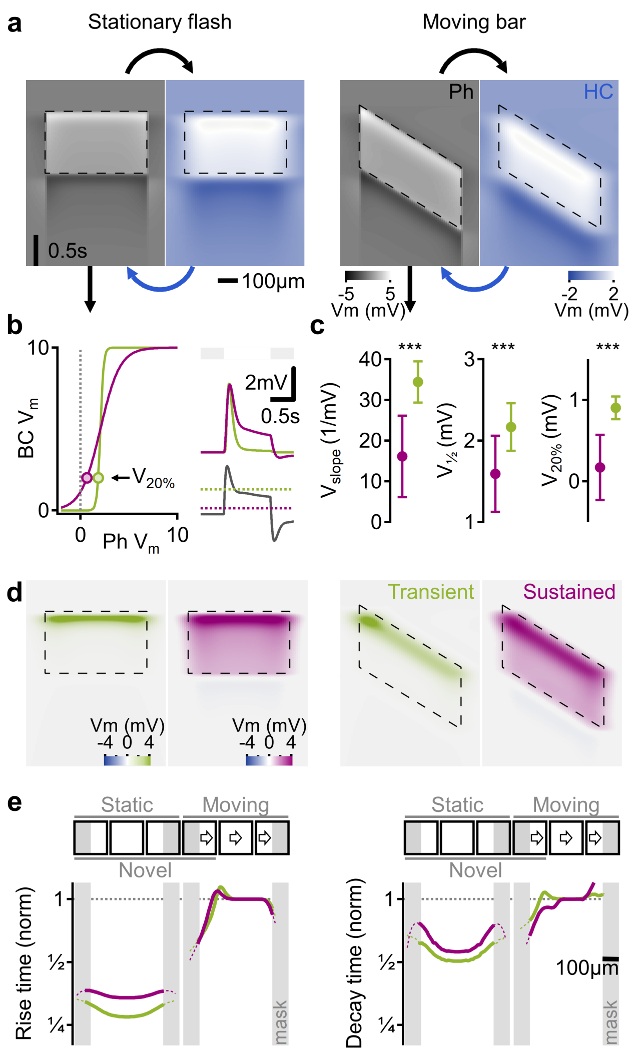


**Fig. S7. Elevated affinity to neurotransmitter release promotes a transient responses in simulated BCs. a** Space-time plot showing the voltage profile of photoreceptors (grey) and HC (blue) in the network simulated in **Fig 3**. **b-c** BC dynamics are influenced by the shape of the signal transformation at the Photoreceptor-BC synapse**. b** Synaptic sensitivity (left) and the simulated light response (right) for two representative BCs (red-sustained, green-transient). V20% marks the photoreceptor voltage that depolarized the BC to 20% of the maximally attainable response (right, bottom, dotted). **c** The effects of the slope (left), half-peak (center) and V20% (right) of the Ph-BC input-output transformation on BC dynamics. ***$p < 0.001$, t-test, corrected for multiple comparisons. Error bars, SD. **d** as in **a** for the two BCs shown in **b**. **e** The temporal for the cells shown in **b** vs. spatial positions. The data is normalized by the full-field motion responses.

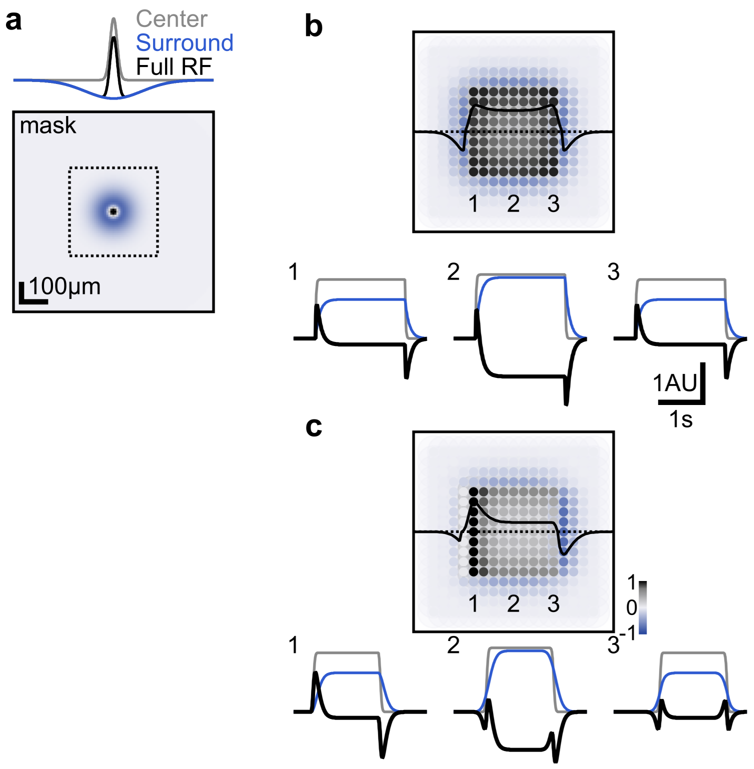


**Fig. S8. Novel object enhancement is a general property of center-surround RFs**

**a** Example spatial extent of a simulated neuron with a linear center-surround RF organization. **b** Top, peak responses from a population of neurons with a similar RF structure to flashed stationary square (stimulus position marked in **a**). The black curve shows a horizontal activation profile of cells near the dotted line. Bottom, the temporal RF evolution in three example cells whose horizontal position is marked on top. Color coding as in **a**. **c** as in (**b**) for a moving bar.

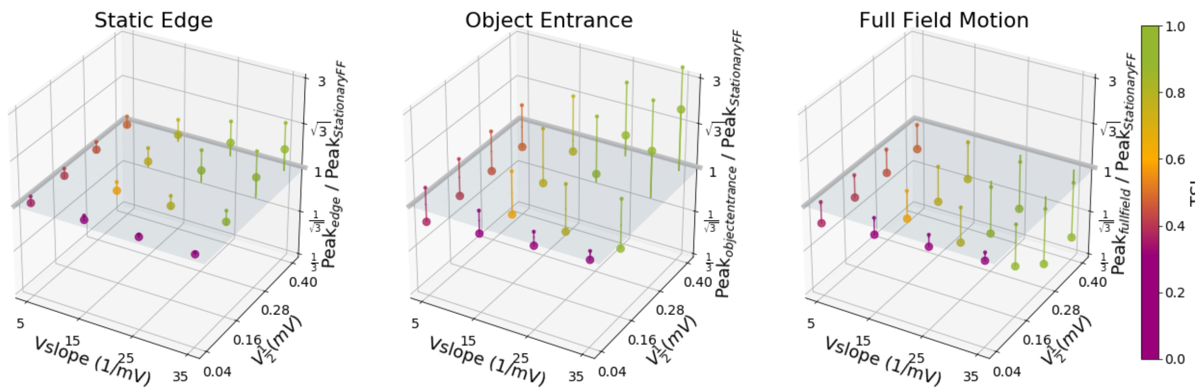


**Fig. S9. Dependence of static and motion processing in BCs on photoreceptor-bipolar synapse parameters**

Simulations were performed to systematically investigate the contribution of the half-width potential, slope, and extent of the bipolar RF on the photoreceptor-bipolar transformation function near a static edge (left), object entrance to the RF from behind a mask (middle), and full-field motion (right). Different parameters were used in each simulation trial and plotted as scatter points in these 3D volumes. The independent variable in these investigations are shown as offsets in the Z dimension, which represent peak response amplitude normalized by the peak stationary full-field response (note the log scale). To indicate Unity (no difference relative to the stationary full-field response) and to aid interpretation, a plane has been plotted at z = 1 and lollipop stalks above or below this plane project to their respective scatter points. Color coding indicates the transiency index computed from the full-field stationary response waveform. Circle sizes indicate the simulated extent of BC RFs, which were 10 μm (small circle) and 60 μm (large circle).

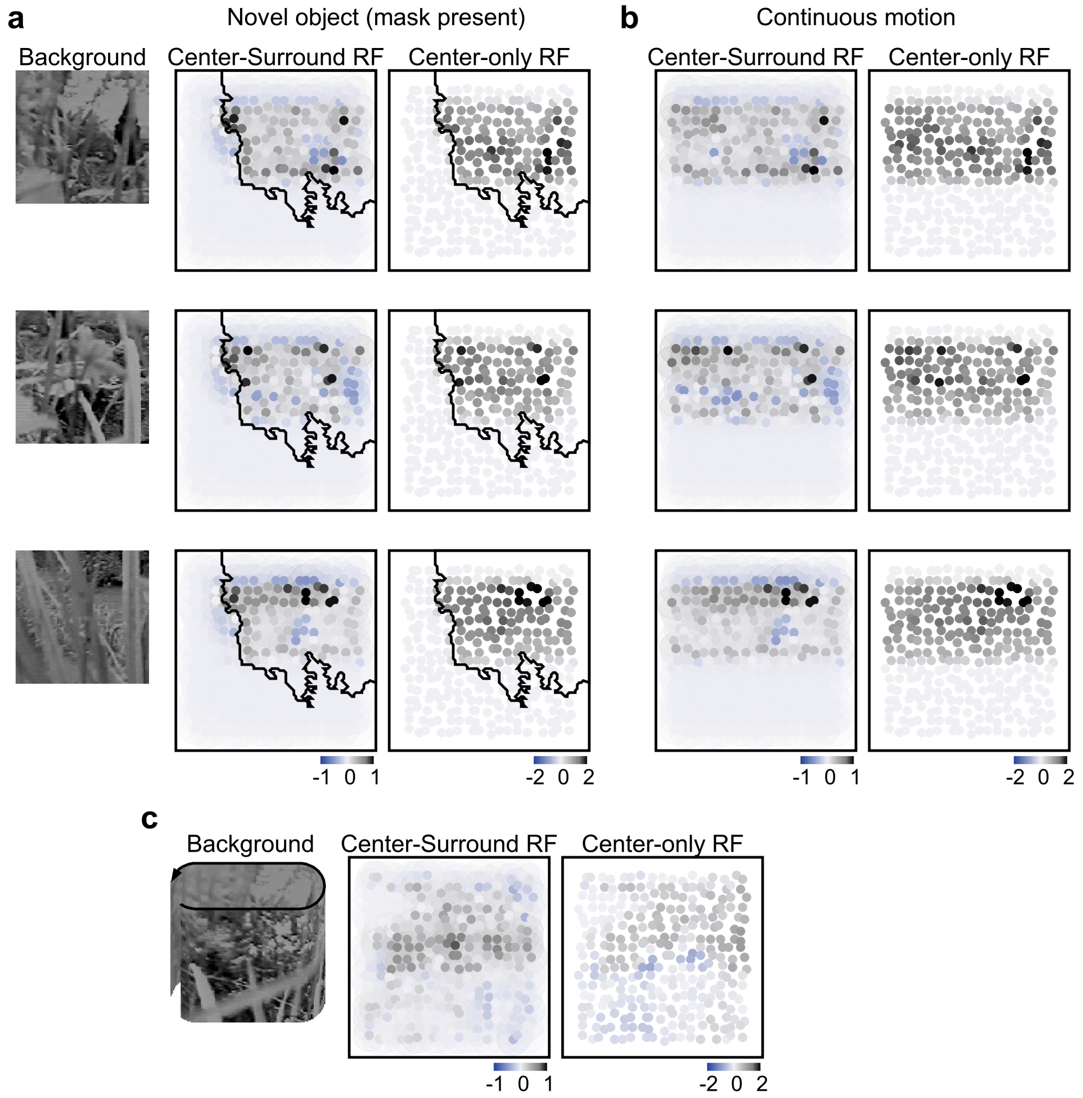


**Fig. S10. Example responses of linear center-surround RFs to natural motion**

**a-b** Example population responses to natural movies with different backgrounds. The stimulus was the predator shown in **Fig. 5,** occluded by a mask (**a**) or moving unoccluded over the entire scene (**b**). **c** Responses to a global, horizontal translation of the background. No stimulus was shown.
